# Targeting tumor stemness switch phenotype by activating pathogen induced stem cell niche defense

**DOI:** 10.1101/2022.03.25.485829

**Authors:** Seema Bhuyan, Bidisha Pal, Lekhika Pathak, Partha Jyoti Saikia, Shirsajit Mitra, Sukanya Gayan, Reza Bayat Mokhtari, Hong Li, Chilakamarti V Ramana, Debabrat Baishya, Bikul Das

## Abstract

Cancer stem cells (CSCs) reside in their tumor microenvironment (TME) niches, which are often hypoxic. Previously, we found that hypoxia and oxidative stress prevalent in TME may reprogram CSCs to a highly aggressive and inflammatory phenotype, the tumor stemness switch (TSS) phenotype. We previously reported a “stem cell niche defense” mechanism in bone marrow and lung mesenchymal stem cell niche against pathogen. Pathogen induced bystander apoptosis (PIBA) of stem cells harboring intracellular pathogen may be part of this defense mechanism. We speculate that the TSS phenotype may also activate this niche defense mechanism to defend their TME niche against pathogen and therefore could be exploited to target CSCs. Here we report that CSCs of TSS phenotype enriched in post-hypoxia ABCG2+ CSCs of several cell lines of diverse tumors including oral squamous cell carcinoma cell line SCC-25 exhibited bystander apoptosis when infected with either Bacillus Calmette Guerin (BCG) or mutant *Mycobacterium tuberculosis (<u>Mtb</u>) strain 18b*. The conditioned media (CM) of the infected cells not only exhibited marked anti-tumor activity in vivo, but also showed significant anti-microbial activity. A detailed mechanisms study revealed that some of the infected ABCG2+ CSCs underwent pyroptosis and released a high mobility group box protein 1 (HMGB1)/p53 death signal that can induce a toll like receptor (TLR) 2/4 mediated bystander apoptosis. Thus, our findings suggest that PIBA can be utilized to activate the “niche defense” mechanism in TSS phenotype, which not only target the TSS, but also exhibit marked anti-tumor activity in vivo.

## Introduction

Cancer stem cells (CSCs) are endowed with self-renewal capacity, and reside in specific niches in the tumor microenvironment (TME). CSCs reside in both the perivascular and hypoxic niches and maintain a steady proportion among the heterogeneous cancer cell population [1]. CSCs are found to express drug efflux pumps such as ATP Binding Cassette Subfamily G Member 2 (ABCG2) [1], exhibit marked detoxification and anti-oxidant activities which makes them inherently resistant to chemo and radiation therapy. Moreover, targeted therapies that inhibit oncogene or growth factor driven pathways fail to target CSCs [2] as these self-renewing cancer cells may activate multiple growth factor pathways. Anti-angiogenesis therapy may aid into the CSC maintenance by increasing tumor hypoxia [3]. Furthermore, immunomodulatory attributes of CSCs make them exceptionally adept at evading immune monitoring [4], and also resistant to immunotherapy. Thus, CSCs are difficult to target by conventional anti-cancer strategies.

Importantly, hypoxia and oxidative stress prevailing in TME may reprogram CSCs to a highly inflammatory and aggressive phenotype, the tumor stemness switch (TSS) phenotype [19]; [52], [53]. Moreover, chemotherapy and radiation induced oxidative stress may also reprogram CSCs to this TSS phenotype [5]. Earlier we reported a cisplatin mediated TSS phenotype in migratory side population (SP) cells; following cisplatin therapy these cells exhibited rapid self-renewal, expressed stemness genes such as octamer-binding transcription factor 4 (Oct-4) and Nanog as well as secreted angiogenic growth factors [5]. Others also reported drug-induced stemness in several tumor models [6], [7]. Subsequently, using a SCC-25 oral cancer derived CSC model, we showed that chemotherapy-induced TSS enable CSCs to modulate TME for rapid tumor progression and immune suppression [8], [9]. In addition, we found that oral CSCs undergo inflammation and bacteria mediated TSS phenotype [10], [11], which also contribute to rapid tumor progression. Therefore, there is a strong rationale to develop innovative therapies to target CSCs exhibiting TSS phenotype.

Apoptosis reinstatement has been a promising anti-cancer strategy; bystander apoptosis incurred by activated macrophages, if suitably courted, could be significant to treatment success [12]. Cancer cells undergoing apoptosis may activate macrophage mediated innate immune mechanism of phagocytosis leading to bystander apoptosis of cancer cells [12]. However, CSCs may suppress the macrophage mediated cancer cell killing [13].

So far, the potential of pathogen induced apoptosis (PIA) in targeting CSCs has not been investigated. PIA is an innate defense system of innate immune cells that has evolved to defend hosts from invading pathogens [14]. When intracellular pathogen such as *Mtb* internalizes in macrophages, these infected cells undergo PIA to confine and restrain the mycobacterial growth [15]. PIA may also involve pyroptosis, a mode of cell death that can spill pathogen-induced molecular pattern (PAMP) and damage associated molecular pattern (DAMP) such as high mobility group box protein 1 (HMGB1) into the microenvironment [16], [27]. Importantly, PIA may involve bystander apoptosis; *Mtb* infected macrophages were found to induce apoptosis in neighbouring uninfected macrophages [15]. Our work on *Mtb* infected MSC model indicates that pathogen induced bystander apoptosis (PIBA) may be part of the stem cell niche defense, an innate defense mechanisms that involves stem cell altruism [25], [40], [53].

We hypothesize that BCG and *Mtb* may cause PIBA of CSCs. Importantly, TSS phenotype of CSCs may activate the “stem cell niche defense” mechanism to defend their TME niche similar to altruistic stem cells (ASCs) [53], [25]. This innate defense mechanism may involve not only secreting anti-microbial factors, but also eliminating self to avoid being attractive site for pathogen’s replication. Hence, we speculate that this PIBA could be utilized to target CSCs of TSS phenotype in the TME niche. Here, we tested this hypothesis by using an invitro model of *Mtb* and *BCG* infected CSCs. BCG is an attractive candidate for PIBA of CSCs, as it is already in clinical use against bladder cancer for the last four decades [17], [18].

We found that *BCG* preferentially infects and replicates intracellular to CSCs of TSS phenotype obtained from diverse cancer cell lines including SCC-25 oral cancer cell line [16], [46]. BCG infected CSCs of TSS phenotype undergo pyroptosis and releases a death signal, the HMGB1/p53 complex, which then induces bystander apoptosis of CSCs in a TLR 2/4 dependent manner. These results identify a novel mechanism of PIA having potential therapeutic implications.

## Results

### BCG replicates intracellular to ABCG2+ CSCs and induces pyroptosis

We hypothesized that BCG and *Mtb* may induce pathogen induced bystander apoptosis (PI-BA) of CSCs exhibiting TSS phenotype, as this phenotype may activate the “stem cell niche defense” mechanism to defend their TME niche similar to ASCs [53], [25]. This innate defense mechanism may involve not only secreting anti-microbial factors, but also eliminating self to avoid being an attractive site for pathogen’s replication. To test this hypothesis, we have utilized several cancer cell line-derived xenograft models, where we characterized the TSS phenotype. In these xenografts of neuroblastoma (SKN-BE-2), sarcoma (HOS and RH4), small cell lung cancer (H-146), colon cancer (LOVO), breast cancer (MCF-7), and oral squamous cell cancer (SCC-25), we found that ABCG2+ CSCs having high tumorigenic activity reside in the hypoxic TME niche and exhibit TSS phenotype [19], [46], [54], [55]. We previously obtained this highly tumorigenic ABCG2+ CSCs from side population (SP) cells [43] when exposed to 24 hours of hypoxia followed by 24 hours of re-oxygenation [19]. We termed these cells as post hypoxia migratory SP cells or SPm (hox) cells [19]. These SPm (hox) cells exhibit TSS phenotype, and enriched in ABCG2+ CSCs [19], [52]. The post hypoxia non-migratory SP cells or SPn (hox) cells were enriched in ABCG2-CSCs [19], [52]. When these cancer cell lines are exposed to an in vitro system of hypoxia and reoxygenation, the ABCG2+ CSCs (equivalent to migratory SP cells) reprogram to TSS phenotype [19], [52]. These post-hypoxia/reoxygenation ABCG2+ CSCs or TSS phenotype, when infected with *Mtb-m18b* or BCG and cultured in vitro for two weeks (Figure 1A), there was 4-5 fold loss of viability as compared to post hypoxia/reoxygenation ABCG2-CSCs (p<0.001, Figure 1B-C), and or pre-hypoxia SP cells. Importantly, the conditioned media (CM) of ABCG2+ CSCs exhibited 3-4 fold anti-microbial toxicity (p<0.05, Figure 1D), suggesting that the TSS may defend their niches from pathogen infection. Our result also suggests that to defend the niche, the TSS also exhibited the PIBA of neighbor ABCG2+ CSCs (Figure 1B-C).

**Figure 1:**
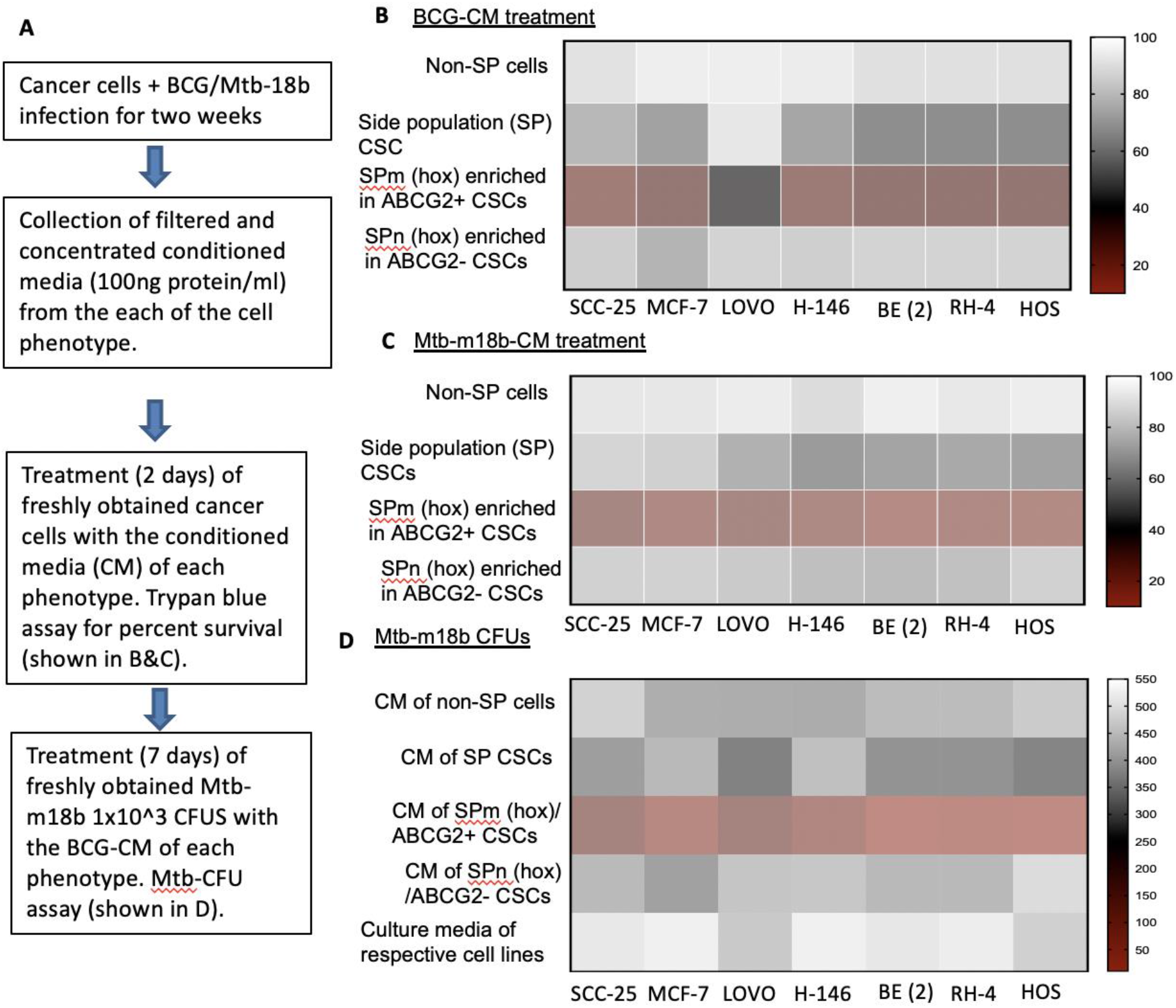
The TSS phenotype of ABCG2+ CSCs exhibit BCG or Mtb-m18b mediated bystander apoptosis and anti-microbial activity. **A**. Experimental plan. To investigate the niche defense potential of each phenotype, we treated the CM of infected phenotype with the untreated corresponding phenotype. **B&C**. Marked bystander cell death is seen in the ABCG+ CSCs group. D. The CM of ABCG2+ CSCs exhibit anti-microbial activity against Mtb-m18b.

We reasoned that exploring the BCG-induced bystander apoptosis in oral cancer CSCs may have clinical utility, as oral cancer lesions are externally accessible for future BCG immunotherapy. Hence, we decided to study the potential PIBA in the SCC-25 cell line. Thus, the ABCG2+ CSCs/ABCG2-CSCs were immunomagnetically sorted from post hypoxia/oxidative stress treated SCC-25 cells [19]. The sorted CSCs were infected with GFP-tagged BCG as previously described [20] and subjected to confocal microscopy. The *Mtb*-colony forming unit (CFU) was performed after 4 days of in vitro cell growth of CSCs infected with green fluorescent protein (GFP)-tagged BCG. *Mtb*-CFU assay confirmed the internalization and replication of the GFP-BCG mostly to ABCG2+ CSCs versus ABCG2-CSCs (Figure 2A & B). The result suggests that the BCG pathogen may selectively infect and replicate in ABCG2+ CSCs versus ABCG2-CSCs. The intracellular replication of pathogen in ABCG2+ CSCs versus ABCG2-CSC is not restricted to *Mycobacterium bovis*, as similar result was observed when the CSCs were infected with an *Mtb* strain m18b, and the infected cells were grown for 4 days [39]. This selective uptake of BCG/*Mtb* by ABCG2+ CSCs, as well as their intracellular replication allows us to evaluate long-term fate of these infected cells.

**Figure 2:**
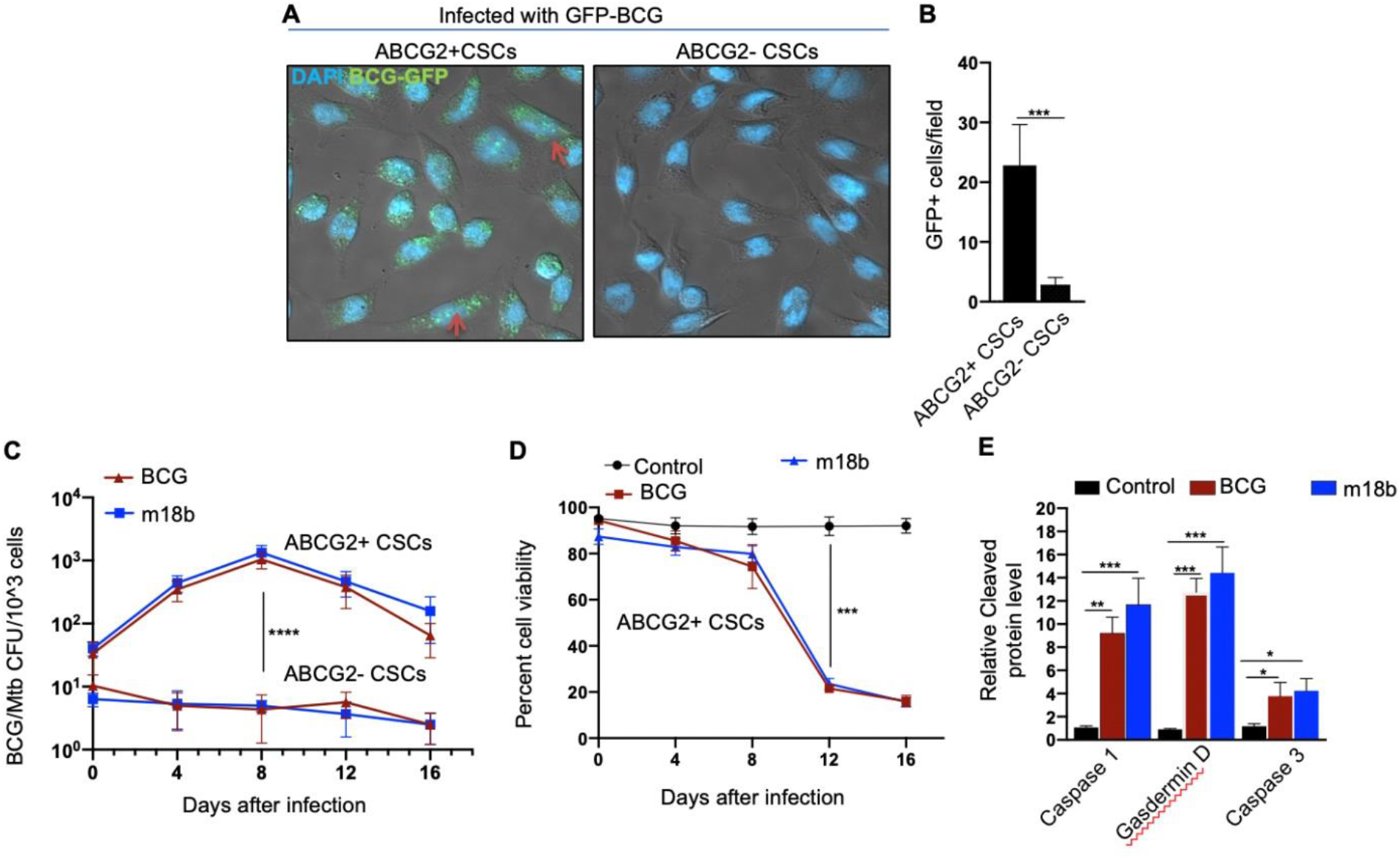
BCG replicates intracellular to ABCG2+CSCs of SCC-25 cell line and induces pyroptosis. ABCG2+ CSCs exposed to the in vitro system of hypoxia and reoxygenation [19] is sensitive to BCG-induced cell death. **A.** Confocal microscopy images (magnification 20x) showing the localization of GFP-positive BCG intracellular to ABCG2+CSCs (shown with arrows) versus ABCG2-CSCs. B. Histogram shows 20-fold increase in GFP-positive BCG per hundred microscopic field in ABCG2+ CSCs versus ABCG2-CSCs. **C.** Intracellular BCG/Mtb m18b-CFU in ABCG2+CSCs versus ABCG2-CSCs at various days after infection. **D.** Trypan blue assay of cell viability of BCG/Mtb-m18b infected ABCG2+CSCs at various days after infection. The control group is the ABCG2+CSCs without any infection **E**. ELISA based measurement of cleaved Caspa-se-1, Gasdermin D and Caspase-3 levels in ABCG2+ CSCs of day-12 after BCG/Mtb-m18b infection. Data represent means ± SEM (B-D). N=3 independent experiments*p<0.05, **p<0.01,*** p<0.001, ****p<0.0001(student t-test).

In macrophages, *Mycobacteria* are known to replicate during the first week of infection, and subsequently, the host cells undergo cell death by apoptosis as well as pyroptosis by the second week of infection [21]. To evaluate whether BCG and *Mtb-m18b* infected ABCG2+ CSCs may also undergo cell death by apoptosis and pyroptosis due to intracellular replication of the pathogen, the infected cancer cells (10^4^/ml) were grown in vitro for 16 days to find out the day, when pathogen replication goes down with associated increase in host cell death. BCG/*Mtb*-*m18b* infected ABCG2-CSCs served as control. Thus, every 4^th^ day, 5x10^3^ cells were recovered, subjected to try-pan blue viability assay, and then lysed to perform the BCG/*Mtb-m18b* CFU assay for evaluating intracellular bacterial replication. Indeed, BCG/*Mtb-*m18b infected ABCG2+ CSCs showed a 100-fold (p<0.0001; Figure 2C) increase in the number of intracellular CFUs on day 8 without exhibiting any marked loss in cell viability (Figure 2D). These results further confirm that the pathogens selectively infect and replicate in ABCG2+ CSCs versus ABCG2-CSCs. Notably, the pathogen infected ABCG2+ CSCs showed marked loss of intracellular CFUs between day 8 and 16 (Figure 2C), as well as 4.5-fold loss of viability between day 8 and 12 (p<0.001; Figure 2D). On day-12, the infected ABCG2+ CSCs exhibited significant up-regulation of caspase-3, as well as caspase-1, a marker of pyroptosis (Figure 2E). These results indicate that the *Mycobacteria* selectively infect, replicate and then induce apoptosis/pyroptosis in ABCG2+ CSCs of SCC-25 cell line.

### The CM of BCG infected ABCG2+ CSCs induces bystander apoptosis in non-infected ABCG2+ CSCs

Next, we evaluated the potential induction of bystander apoptosis by the CM of BCG and *Mtb-m18b* infected ABCG2+ CSCs. On day 12, the CM of BCG or *Mtb-m18b* infected ABCG2+ CSCs was collected, filter sterilized with 0.2µm filter, concentrated with Centricon centrifugal filter units (EMD Millipore) to 10X and then used to treat fresh ABCG2+ CSCs. The CM treated cells were evaluated for the *Mtb*-CFUs, cell viability and caspase 1 & 3 protein levels. The 72 hours CM treated fresh ABCG2+ CSCs showed 4-5 fold reduction in cell viability (p< 0.0001; figure 3A), without any evidence of *Mtb*-CFU growth. The loss of cell viability was associated with 10-12 fold increase in cleaved caspase 3 level whereas, the level of cleaved caspase-1 remained unchanged (Figure 3B). These results suggested the induction of bystander apoptosis by the CM of BCG as well as *Mtb-m18b* infected ABCG2+ CSCs.

**Figure 3:**
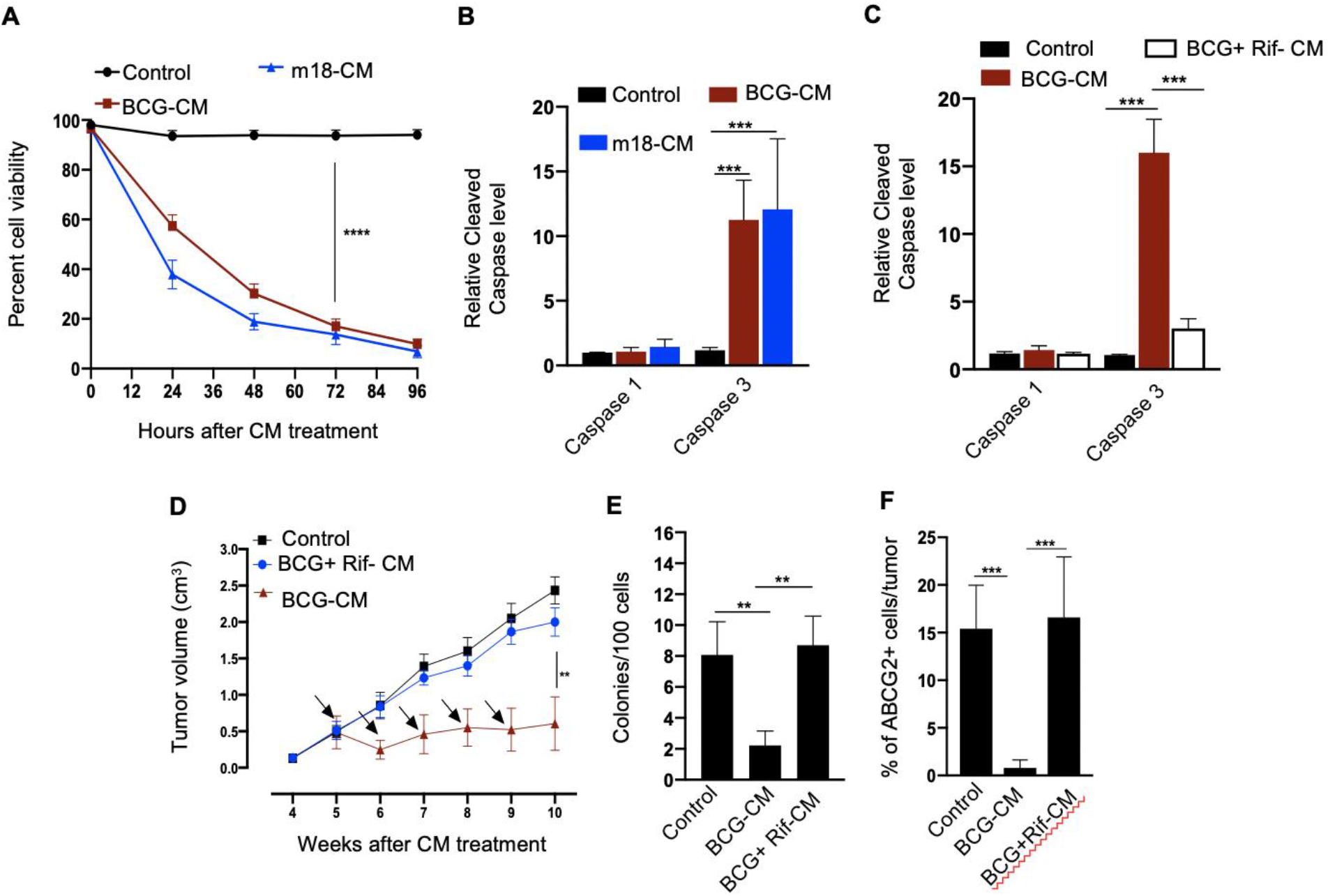
The CM of BCG infected ABCG2+CSCs induces bystander apoptosis in non-infected ABCG2+ CSCs. **A**. Trypan blue assay of cell viability of non-infected ABCG2+CSCs after treat-ment with CM of BCG/Mtb-m18b infected ABCG2+ CSCs. The cell count was performed at various hours after the treatment. B&C. ELISA results of cleaved Caspase levels following 48 hours of treatment with the CM of BCG/Mtb-m18b infected ABCG2+ CSCs. Rif+CM denote the CM of BCG infected ABCG2+ CSCs treated with Rifampicin (Rif). **D.** In vivo growth of tumor in a SCC-25 derived xenograft model of NOD/SCID mice (n=10 in each group) treated with CM of BCG infected ABCG2+CSCs. The tumor volume was measured at various weeks after the CM treatment. Arrow indicates the intra-tumor treatment with 0.1 ml of sterile concentrated CM containing 0.5 mg protein. **E &F.** The clonogenic potential and percentage of ABCG2+ CSC population in dissociated tumor cells obtained from the xenografts of 10^th^ week after the CM treatment. Data represent means ± SEM (B-D). N=3 independent experiments (A-C); N=5 independent experiments (E-F)**p<0.01, *** p<0.001, ****p<0.0001 (t-test). BCG-CM: The CM of BCG infected ABCG2+CSCs, m18b-CM: The CM of Mtb-m18b infected ABCG2+ CSCs, Rif-CM: The CM of Rifampicin treated BCG infected ABCG2+CSCs, Control: The CM of ABCG2+ CSCs not infected with BCG/Mtb-m18b

As per our hypothesis, bystander apoptosis is the result of alarm signals released by the host cells, where pathogen replicates. Therefore, inhibition of pathogen replication in the host cells may significantly reduce bystander apoptosis of neighboring cells. To test this hypothesis, the BCG-infected ABCG2+ CSCs were treated with rifampicin (2µg/ml) for 2 days that kills intracellular BCG [39]. On day-12, the CM (henceforth known as BCG+Rif-CM) was collected, filter sterilized, and added to freshly grown ABCG2+ CSCs. The BCG+Rif-CM treated cells were then evaluated for cell viability, cleaved caspase 1 and 3 levels. BCG-CM treated ABCG2+ CSCs were used as positive control. We found that the BCG+Rif-CM treatment had no significant effect on cell viability and apoptosis (Figure 3C). These results indicate that the replication of BCG intracellular to ABCG2+ CSCs is required for the CM of these cells to induce bystander apoptosis.

Next, we evaluated the in vivo potency of bystander apoptosis of ABCG2+ CSC in a SCC-25 derived xenograft model of NOD/SCID mice, which we recently characterized [8], [41]. The BCG-CM, BCG+Rif-CM or saline (1 ml/week/i.p.) were injected to SCC-25 tumor bearing NOD/SCID mice (n=10 in each group; initial tumor size of 0.5 mm3; Figure 3D). Tumor growth was measured weekly until the control tumor (the group with saline alone treatment) reach maximum size of 2cc (Figure 3D). At the end of the treatment, tumors were dissociated to obtain single cell suspension; the cells were subjected to clonogenic assay, as well as immunomagnetic sorting to obtain the ABCG2+ sub-population cells, and their cleaved caspase 3 levels. Results are given in Figure 3 D-F. We found a 4-fold decrease in the tumor volume after 5 weeks of treatment in the BCG-CM versus BCG+Rif-CM treated group (Figure 3D-E). Importantly, in the clonogenic assay, the ABCG2+ subpopulation exhibited a 15-fold reduction in the BCG-CM treated group (Figure 3 F). Due to this low number of ABCG2+ CSCs, we could not measure the caspase 3 level in these cells. Nevertheless, these results indicate the ability of the BCG-CM to target the CSC population, as well as reduce the tumor growth in mice.

### Intrinsic apoptotic pathway is involved in bystander apoptosis of ABCG2+ CSCs

Next, we investigated the cellular and molecular mechanisms of BCG-CM induced bystander apoptosis. We have considered that the BCG infection induced pyroptosis may have released putative alarm signals capable of inducing bystander apoptosis in neighboring CSCs (Figure 4A). Indeed, BCG infection was associated with lactate dehydrogenase (LDH) release by ABCG2+ CSCs (Figure 4B). Treatment of the BCG infected cells with disulfiram (50nM/twice daily for 4 days starting on day 8), or Capase -1 inhibitor, which are inhibitor of pyroptosis, led to a marked reduction in LDH release (Figure 4B). Importantly, the BCG-CM collected from the disulfiram treated cells showed marked reduction in bystander apoptosis (Figure 4C&D), further suggesting that pyroptosis played a significant role in bystander apoptosis by releasing soluble factors.

**Figure 4:**
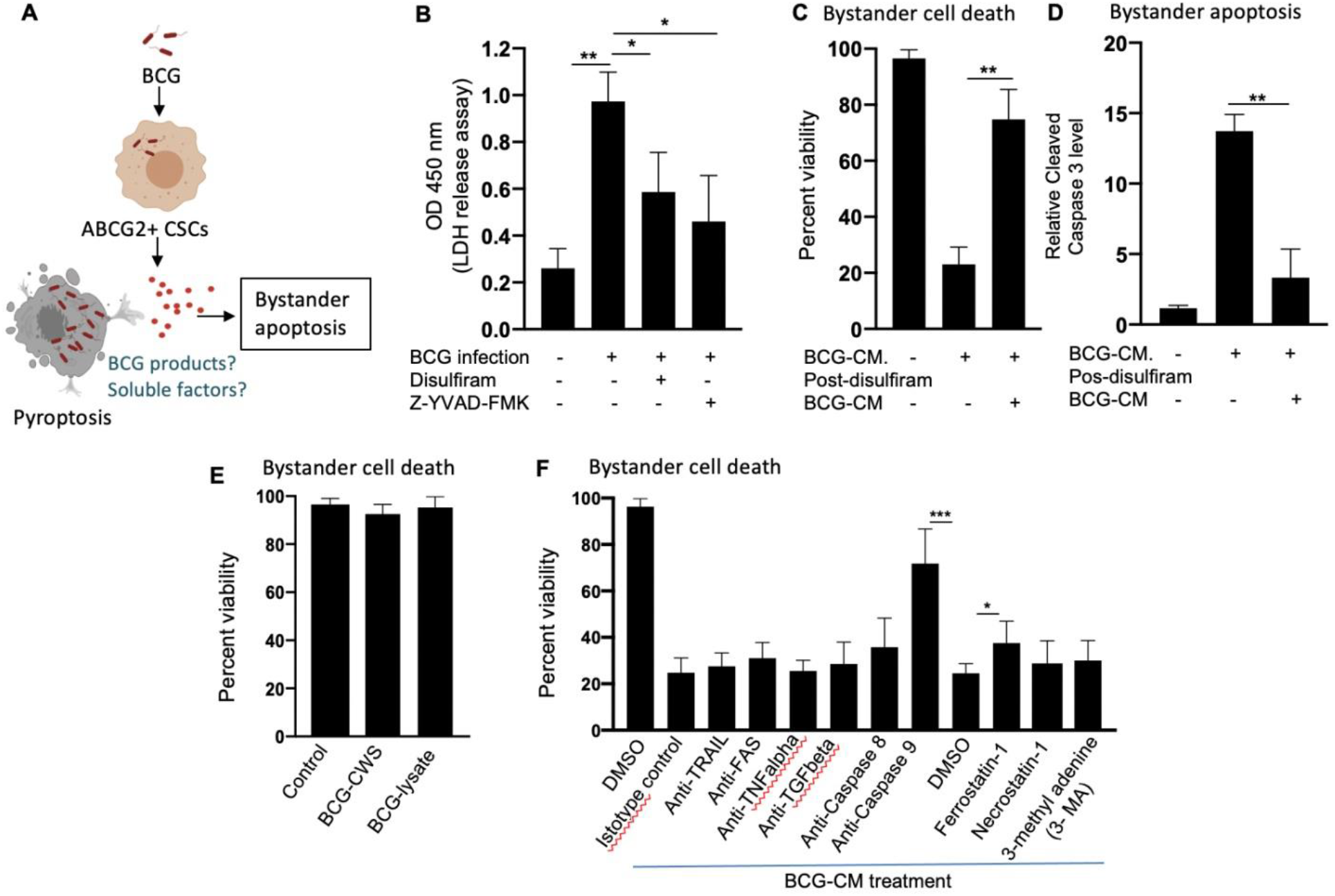
Pyroptosis mediated secretion of soluble factors may induces bystander apoptosis. **A**. The schematic is showing the experimental hypothesis. **B**. The histogram is showing LDH release by BCG infected ABCG2+ CSCs on day-12. The LDH was measured after treating the BCG-infected ABCG2+ CSCs with or without disulfiram and Z-YVAD-FMK from day 8-12. **C&D**. The data is showing uninfected ABCG2+ CSC viability and bystander apoptosis following treatment of BCG-CM obtained from the infected ABCG2+ CSCs with or without disulfiram treatment **E**. ABCG2+ CSCs are not sensitive to BCG cell wall skeleton (BCG-CWS) and BCG-lysate treatment for a week. **F**. ABCG2+ CSCs were treated with various antibodies and inhibitors during BCG-CM treatment. N=3 independent experiments for B-D, and n=4 independent experiment for E&F, One-Way ANOVA (4B), students t-test (4C, D, F). *p<0.05, **p<0.01,*** p<0.001 (student t-test).

We therefore considered identifying the soluble factors released by pyroptotic cells involved in bystander apoptosis. We first speculated that BCG cell wall skeleton (BCG-CWS), and or BCG-derived DNA/RNA may be released by pyroptotic cells, thus present in the BCG-CM, and mediate bystander apoptosis. Previously, these BCG derived products (BCG-CWS or BCG derived DNA/RNA) were found to induce apoptosis in bladder cancer cells by activating TLR 2, 4, 7 and 9 and thus inducing the Myeloid differentiation primary response gene 88 (MYD88) pathway of extrinsic apoptosis [22], [23]. However, BCG-CWS [23], as well as the live-BCG lysate (1x10^7 BCG in 1 ml of DMEM incubated for 24 hours, and then sterile filtered) failed to induce bystander apoptosis in ABCG2+ CSCs (Figure 4E).

Next, we considered the potential release of soluble factors such as TNF-related apoptosis-inducing ligand (TRAIL), FAS ligand, tumour Necrosis Factor alpha (TNF-alpha), and transforming growth factor beta (TGF-beta) in the BCG-CM. These soluble factors may induce apoptosis by activating caspase-8 mediated extrinsic apoptosis [49]. Previously, Kelly DM et al, while studying the mechanism of BCG-induced bystander apoptosis in macrophages and T-cells did not identify soluble alarm signals for bystander apoptosis [15]. Similarly, we also did not find any significant role of these soluble factors in bystander apoptosis; treating the BCG-CM with neutralizing antibodies of these soluble factors did not reduce bystander cell death (Figure 4F). Moreover, inhibition of caspase-8 did not affect bystander apoptosis (Figure 4F), suggesting that extrinsic pathway was not involved in bystander apoptosis. Thus, it is unlikely that soluble factor mediated extrinsic apoptosis was involved in BCG-CM mediated bystander apoptosis.

In contrast, inhibition of intrinsic apoptosis, which is caused by caspase-9 [50] markedly, reduced bystander apoptosis (Figure 4F). Thus, it appears that BCG-CM may contain soluble factors that may internalize into the target cancer cells to induce intrinsic apoptosis. We also considered the potential involvement of other mode of cell death including necroptosis, ferroptosis and autophagy in bystander apoptosis by pretreating ABCG2+ CSCs with a RIP1-kinase inhibitor (Necrostatin-1), ferroptosis inhibitor (Ferrostatin-1) and an autophagy inhibitor (3-methyladenine; 3 MA) before treatment with BCG-CM. Relative to vehicle control (DMSO), Ferrostatin -1 prevented the effect of BCG-CM induced loss of cell viability (Figure 4F). Conversely, the phenotype was not reversed by Necrostatin-1 or 3MA, indicating that necroptosis or autophagy does not affect the ability of BCG-CM to induce death of ABCG2+ CSCs. Taken together, BCG-CM may contain soluble factors that internalize into ABCG2+ CSCs to activate intrinsic apoptosis pathways.

### Bystander apoptosis of ABCG2+CSCs is associated with HMGB1/p53 death signal

Previously, we identified an intrinsic pathway of apoptosis mediated by p53/MDM2 oscillation [24] and HMGB1 [54] in the ASC phenotype [24], [25]. HMGB1, a DAMP associated with alarm signaling of innate defense [54] is actively secreted by cancer cells during stress including hypoxia [47]. Moreover, HMGB1 is secreted by BCG infected immune cells. The extracellular HMGB1 regulates inflammation and play a pro-tumorigenic role [47]. Therefore, it is unlikely that HMGB1 alone would induce bystander apoptosis of CSCs. Previously, it was found that in a colon cancer cell line, HMGB1 binding to p53, along with reactive oxygen species (ROS) production may induce apoptosis and autophagy [26]. Nuclear Magnetic Resonance (NMR) spectroscopy as well as in silicon protein structural analysis indicate that p53 can bind to HMGB1 to make stable HMGB1/p53 complex [30]. Thus, it is possible that pyroptotic cells may release death signal, the HMGB1/p53 complex that mediate bystander apoptosis of neighbor CSCs in the TME.

To investigate this possibility, we did a series of experiments. We first measured HMGB1 and p53 protein levels in the BCG-CM by enzyme linked immunosorbent assay (ELISA) using iMark Microplate Absorbance Reader (Biorad, Gurgaon, India) as well as western blot (WB) assay. Next, we performed co-immuno-precipitation (IP) of p53 and HMGB1 to identify the putative HMGB1/p53 complex in the BCG-CM. The CM of freshly obtained un-infected ABCG2+ CSCs served as a control. The results are given in Figure 5A&C, showing that p53 protein could be detected in the BCG-CM only. Whereas, HMGB1 could be detected in the CM of both BCG infected and also the control group (Figure 5A). However, the IP-WB result clearly demonstrates that only the BCG-CM showed the presence of a HMGB1/p53 complex, as the IP product of HMGB1 contained p53 (Figure 5B). Second, we confirmed that pyroptosis is involved in the secretion of this soluble complex in the CM, as pre-treatment with disulfiram significantly inhibited the secretion of HMGB1/p53 complex by the BCG infected ABCG2+ CSCs (Figure 5C). Third, we performed a protein uptake assay to quantify the potential uptake of the HMGB1/p53 complex by the BCG-CM treated ABCG2+ CSCs. The ABCG2-CSCs served as control. Briefly, p53 was measured in the CM of these cells after they were treated with BCG-CM. Reduction of the p53 concentration in the CM will indicate uptake of this protein by the cells. In this manner, we found that within 4 hours of treatment, ABCG2+ CSCs took 50% of p53 from the BCG-CM (Figure 6A), whereas ABCG2-CSCs took only 6.5%. Thus, there is a 7.5-fold increase of uptake of p53 by ABCG2+ CSCs compared to ABCG2-CSCs (Figure 6B). There was a corresponding decrease of HMGB1 concentration in the CM (11.2 +/- 4.3 ng/ml to 8.4 +/-3.2 ng/ml; p= 0.043, n=5), suggesting the uptake of HMGB1 bound p53 by the ABCG2+ CSCs from the CM. Pre-treatment of BCG-CM with a neutralizing antibody against HMGB1 significantly reduced the p53 uptake by the treated cells (Figure 6A-B), suggesting that ABCG2+ CSCs endocytose HMGB1 bound p53. Fourth, the p53 protein uptake was associated with the induction of p53/mouse double minute 2 homolog (MDM2) oscillation and corresponding activation of p53 down-regulating genes as well as increased level of cleaved caspase 3 in the BCG-CM treated ABCG2+ CSCs (Figure 6C-E). Finally, inhibition of p53 by small molecular inhibitor pifithrin alpha (Figure 6E) or siRNA gene silencing (data not shown) without inhibiting the HMGB1 significantly reduced cleaved caspase 3 level. We found similar result when the HMGB1 was neutralized in the BCG-CM treated ABCG2+ CSCs without inhibiting the p53 (Figure 6E). These findings suggest that the soluble factor HMGB1/p53 complex present in the BCG-CM may be associated with the bystander apoptosis in ABCG2+ CSCs.

**Figure 5:**
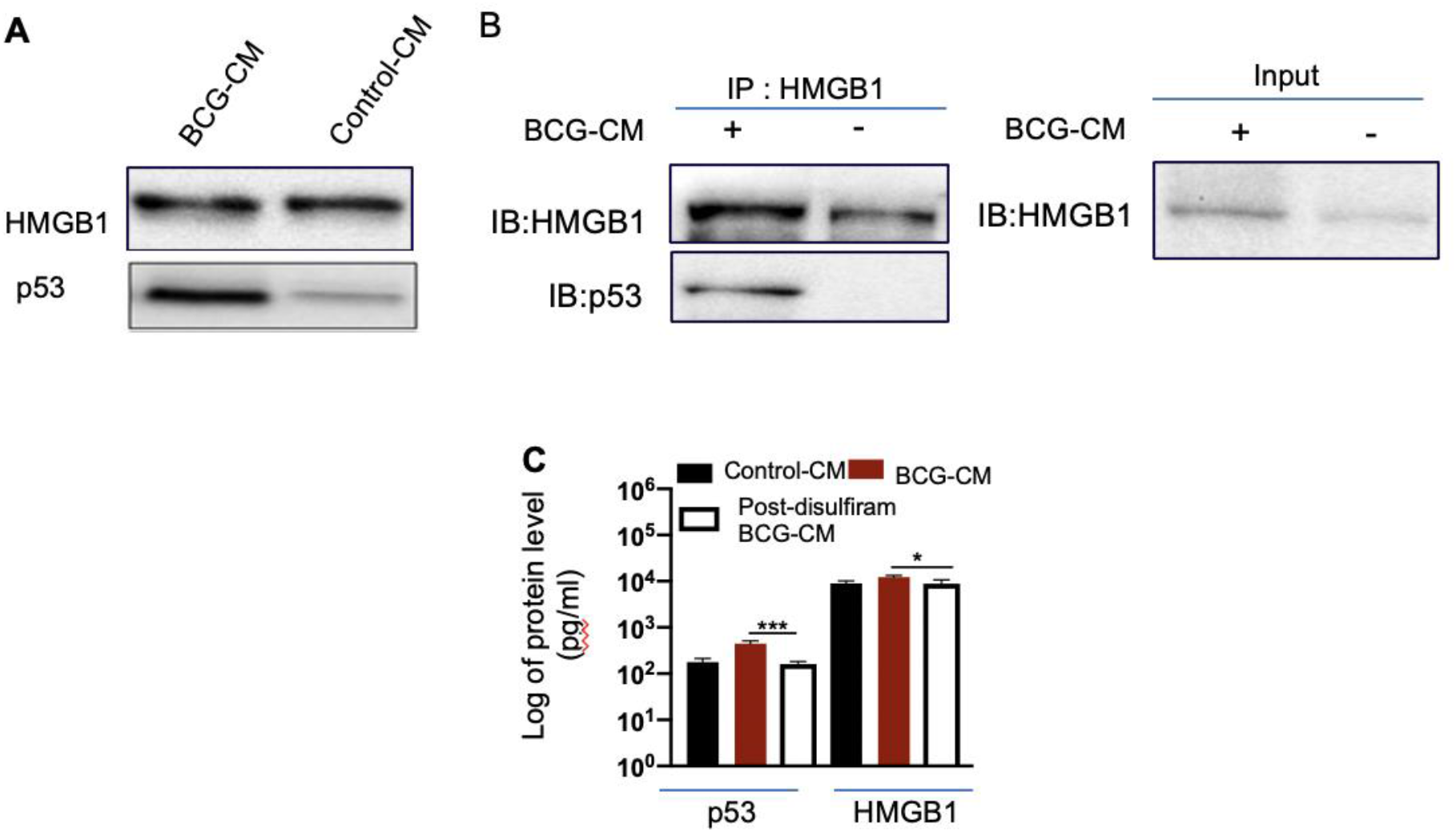
BCG infected ABCG2+CSCs release HMGB1/p53 complex during pyroptosis. **A**. Western blot of concentrated BCG-CM and control-CM showing the presence of HMGB1 and p53. 10 µg protein was loaded in both infected and control group. **B.** Immunoprecipitation experiment confirms the formation of HMGB1/p53 complex in BCG-CM versus Control-CM. Immunoblotting (IB) of HMGB1 and p53 was also performed. Input is 2.5% of the total amount of immunoprecipitated. **C**. The histogram is showing the secretion of HMGB1/p53 complex by BCG-infected ABCG2+ CSCs with or without disulfiram treatment (50nM/twice daily for 4 days). The elute of IP/HMGB1 shown in B was subjected to ELISA, and protein levels were compared with uninfected ABCG2+ CSCs (p53, average 0.13 ng/ml; HMGB1, average 10.5 ng/ml) to obtain fold change. Data represent means ± SEM (A-E). N=3 independent experiments (A-E). *p<0.05, *** p<0.001 (student t-test).

**Figure 6:**
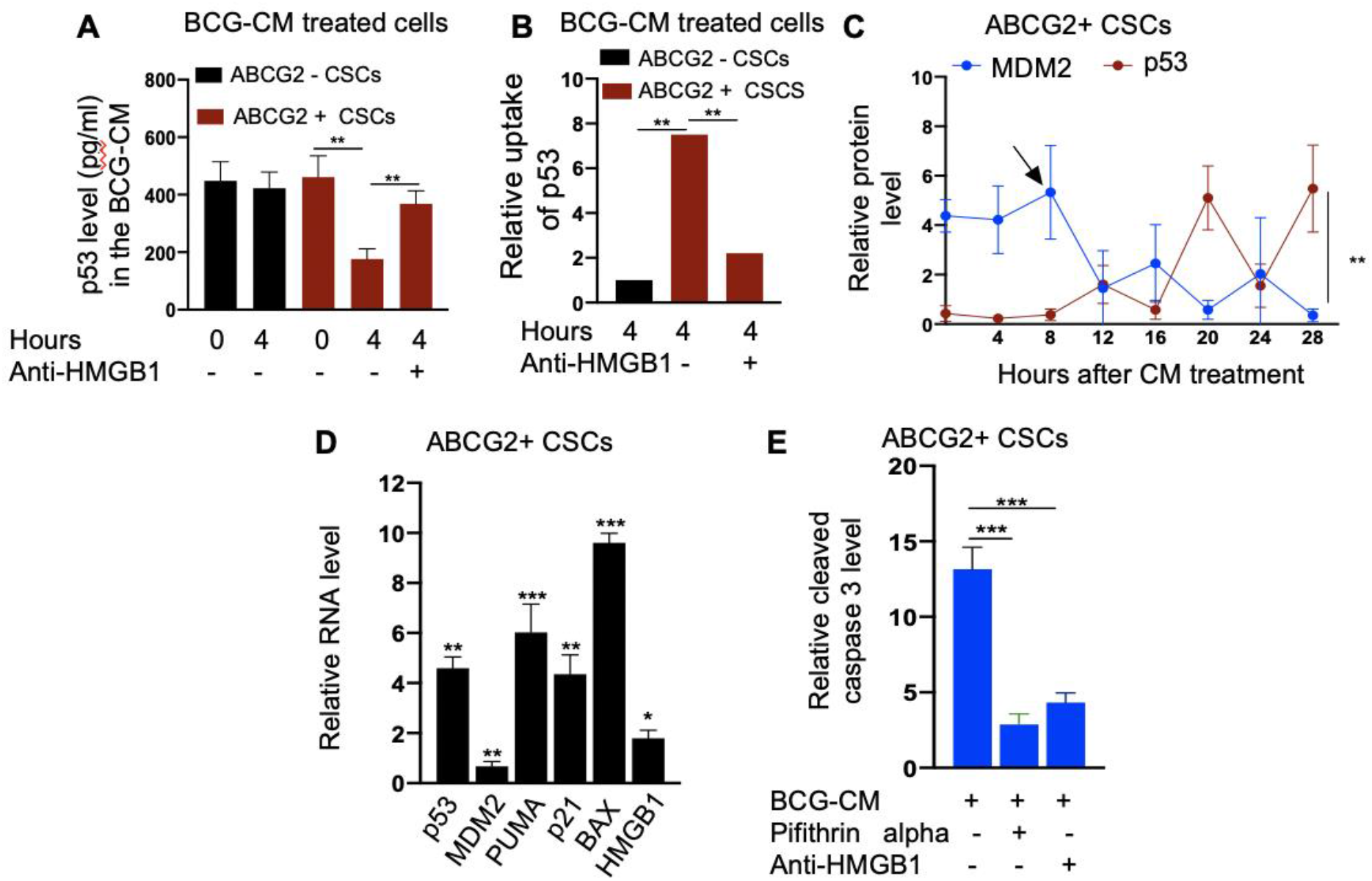
Bystander apoptosis is characterized by the HMGB1/p53 complex mediated apoptosis. **A.** The histogram is showing p53 uptake by the ABCG2+ CSCs and ABCG2-CSCs with or without anti-HMGB1 after 4 hours of treatment with BCG-CM. The p53 was measured in the CM by ELISA. **B.** Protein uptake assay result: the relative uptake of p53 is measured in ABCG2+ CSCs versus ABCG2-CSCs. The absolute uptake of p53 by ABCG2+ CSCs was compared with that of ABCG2-CSCs to obtain fold-change. **C**. The data shows return of p53/MDM2 oscillation in non-infected ABCG2+ CSCs after 12 hours of BCG-CM treatment. The fold change in the protein levels of p53 and MDM2 (measured by ELISA) represents p53/MDM2 oscillation. **D.** Histogram is showing the induction of p53 related apoptotic genes as well as HMGB1 gene in ABCG2+ CSCs following 28 hours of treatment with BCG-CM. The real-time PCR data were compared with untreated ABCG2+ CSCs to obtain fold change. **E**. The histogram shows cleaved caspase 3 level (measured by ELISA) in ABCG2+ CSCs treated with BCG-CM with or without pifithrin alpha (2 µM in DMSO for 48 hours) or anti-HMGB1 (10ug/ml for 48 hours; isotype control of same dose). Data represent means ± SEM (A-E). N=3 independent experiments (A-E). *p<0.05, **p<0.01, *** p<0.001 (student t-test).

### Toll like receptor 2 (TLR 2) and 4 are involved in HMGB1/p53 complex mediated bystander apoptosis

Next, we examined the potential mechanism of the HMGB1/p53 molecular complex uptake into the ABCG2+ CSCs for the induction of bystander apoptosis. Considering that TLR 2 and 4 are well known receptor for exogenous HMGB1, we reasoned that these two receptors may endocytose the HMGB1/p53 complex into the cells (Figure 7A) [47]. Indeed, neutralizing antibodies against TLR 2 and TLR 4, but not TLR 7 and TLR 9 (5µg/ml; Invivogen) significantly inhibited the BCG-CM mediated apoptosis of ABCG2+ CSCs in vitro (Figure 7B) and in vivo (Figure 7C). Moreover, both confocal imaging and biochemical assay showed a marked reduction in caspase 3 expression/activity in ABCG2+ CSCs treated with neutralizing antibody against TLR 2/4 (Figure 7D-F). These data suggest that TLR 2/4 may be involved in BCG-CM mediated bystander apoptosis.

**Figure 7:**
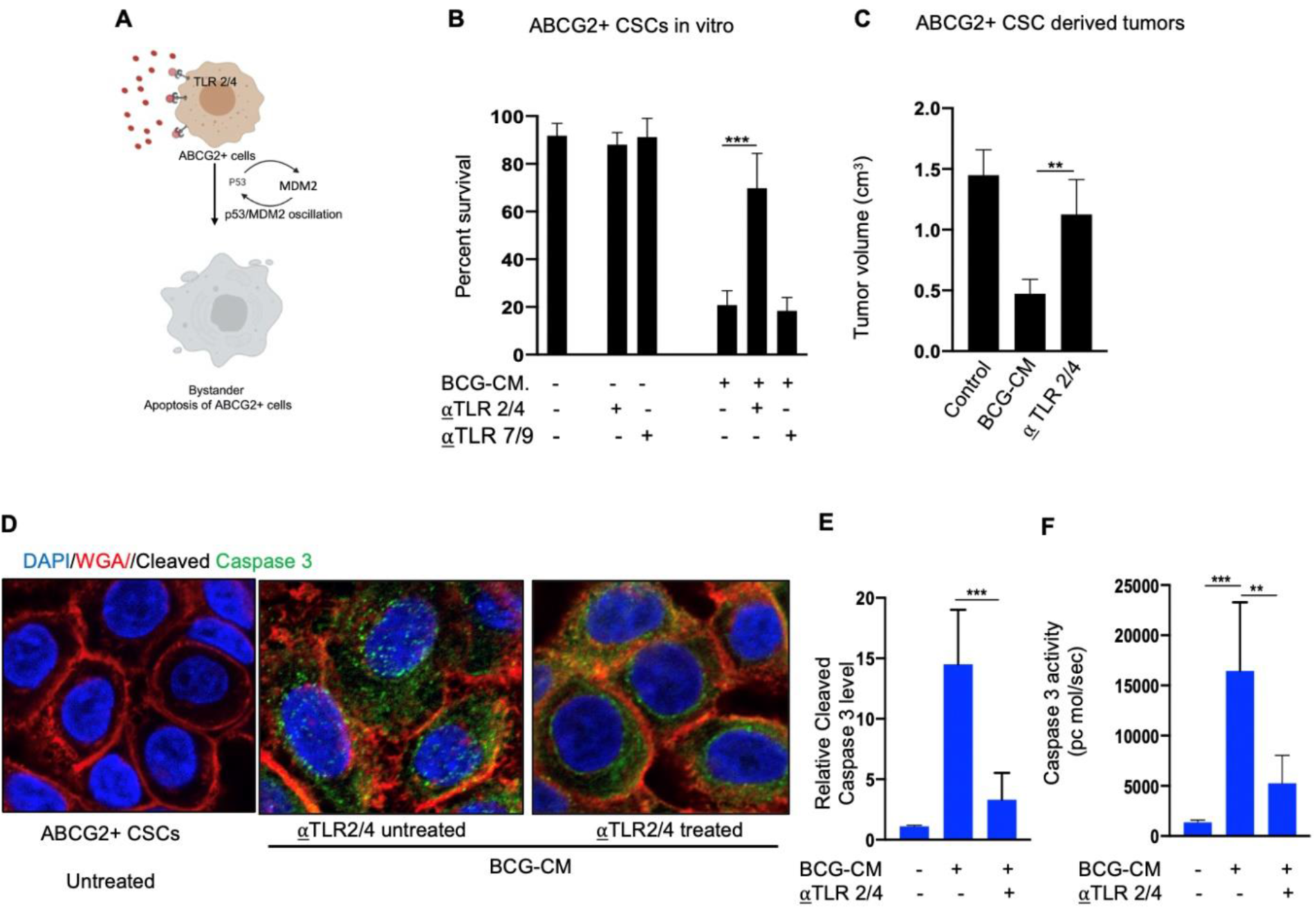
Bystander apoptosis is mediated by TLR 2 and 4. **A.** Hypothesis: TLR 2/4 are required for the execution of HMGB1/p53 complex mediated bystander apoptosis. **B.** Relative cell viability of BCG-CM treated (72 hours) ABCG2+ CSCs pretreated with TLR neutralizing antibodies. C. The histogram is showing the tumor volume in BCG-CM treated and TLR neutralizing Abs pretreated mice group. **D.** Immunofluorescence labeling of apoptotic ABCG2+ CSCs is showing reduction of cleaved caspase 3 staining in the cells pre-treated with TLR 2/4 neutralizing antibodies (Dapi, nuclear stain; WGA, cell membrane stain). Magnification 20x. **E&F**. The histograms are showing corresponding protein level (ELISA) and enzymatic activity of Caspase-3. Data represents means ± SEM (B &D-E). ** p<0.001, *** p<0.0001; N=3, student’s t test.

To further investigate the role of TLR2/4 in the uptake of HMGB1/p53 complex, we revisited the initial finding that ABCG2-CSCs are less sensitive to BCG-CM mediated bystander apoptosis than ABCG2+ CSCs (Figure 1 A and Figure 8A). We reasoned that ABCG2-CSCs may express low level of TLR 2/4 compared to ABCG2+ CSCs leading to the less uptake of the HMGB1/p53 complex. As expected, ABCG2-CSCs expressed 4-6-fold lower level of TLR 2 and TLR 4 gene as well as protein expression (Figure 8B-C). However, there is no significant difference in expression of TLR7 and TLR 9 in ABCG2+ CSCs vs ABCG2-CSCs. Over-expression of TLR2/4 (Figure 8D) led to 2-fold increase in bystander apoptosis and associated 6-fold increase in the uptake of HMGB1/p53 complex (Figure 8E-F). Additionally, this TLR2/4 dependent increase in bystander apoptosis is markedly reduced following inhibition of p53, or neutralizing of HMGB1 in the BCG-CM (Figure 8E-F). These results indicate that TLR 2 and TLR 4 are involved in the HMGB1/p53 complex mediated bystander apoptosis.

**Figure 8:**
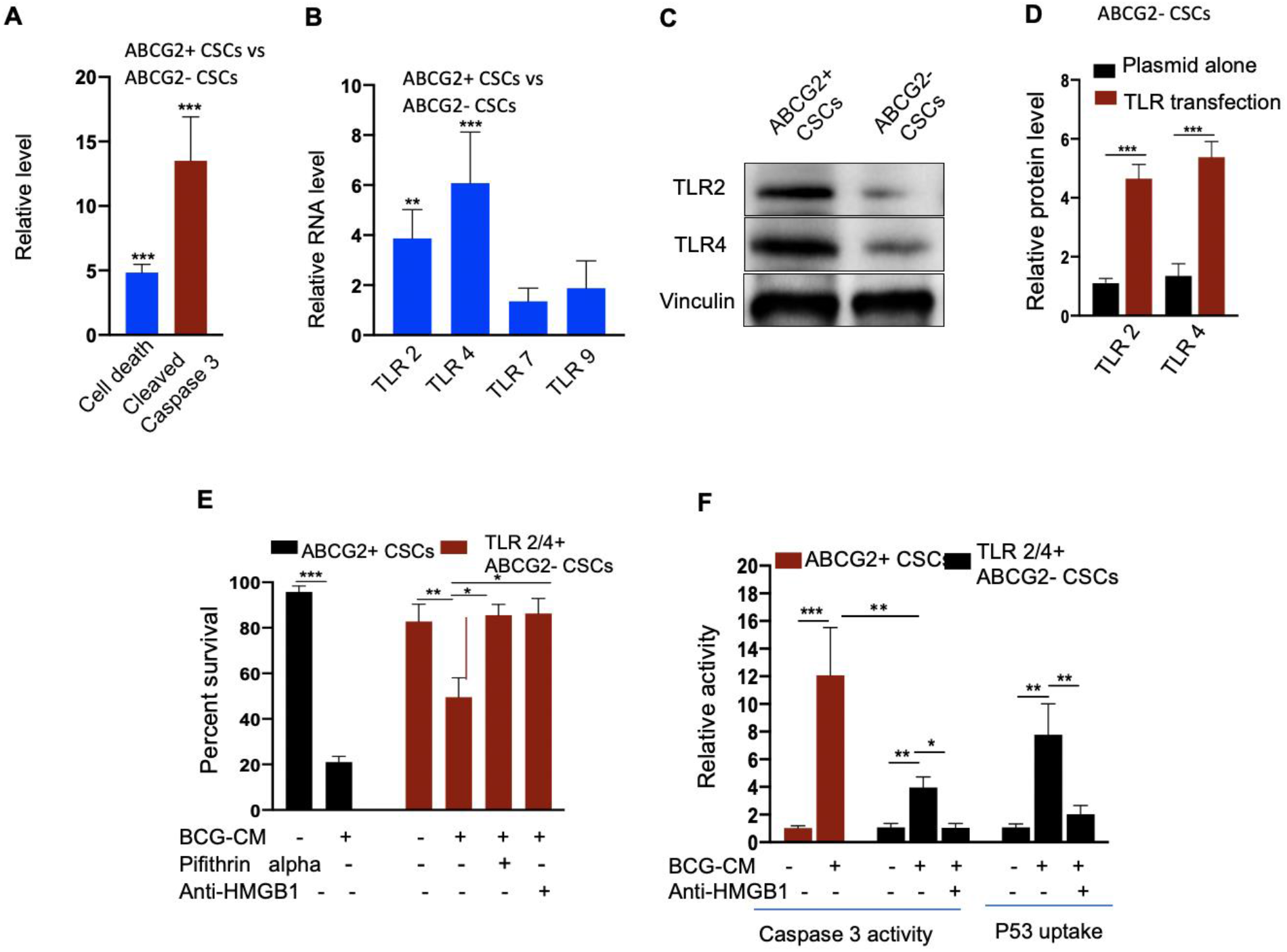
A. TLR 2 and 4 are required for the internalization of HMGB1/p53 complex into the CSCs. **A.** ABCG2-CSCs are significantly less susceptible to BCG-CM mediated cell death and by-stander apoptosis. **B.** Real time PCR data shows fold change in RNA expression of TLR 2, 4, 7 and 9 in ABCG2+CSCs vs. ABCG2-CSCs. **C.** Western blot shows TLR 2/4 expression in ABCG2+CSCs vs. ABCG2-CSCs. **D.** TLR 2 and 4 expressions in ABCG2-CSCs as measured by ELISA. **E.** TLR 2 & 4 overexpressing ABCG2-CSCs show BCG-CM mediated bystander apoptosis that can be reduced by inhibiting p53, and or neutralizing HMGB1 activity. ABCG2+ CSCs served as control for BCG-CM potency. **F.** The Caspase 3 activity was measured after 48 hours whereas p53 uptake activity was measured after 4 hours of BCG-CM treatment. ABCG2+ CSCs served as control for BCG-CM potency. *p<0.05, ** p<0.001, *** p<0.0001, N=3, student’s t test (8A, B, D, F), One-Way ANOVA (8E-F).

### TSS phenotype amplifies the BCG-CM mediated bystander apoptosis

We noted that bystander apoptosis was significantly less in the TLR2 overexpressed ABCG2-CSCs than the ABCG2+ CSCs (Figure 8F), although the HMGB1/p53 (present in the BCG-CM) uptake was similar (Figure 6B and Figure 8F). We hypothesize that in the ABCG2+ CSCs which are of TSS phenotype, the HMGB1/p53 apoptotic signal may be amplified as a part of the CSC niche defense mechanism. Thus, we expect that the p53 concentration in the CM of ABCG2+ CSCs will increase after initial decline. Whereas in the SP cells or and TLR2/4 overexpressing ABCG2-CSCs which do not exhibit TSS phenotype, the death signal would not be amplified. Indeed, we found that the p53 concentration in the ABCG2+ CSCs exhibited a sharp increase by 2.5-fold between 8-16 hours of BCG-CM treatment after initial decline in 4 hours, suggesting the release of fresh HMGB1/p53 complex by the apoptotic cells. Whereas, the p53 concentration in the culture supernatant of SP cells and TLR2/4 overexpressing ABCG2-CSCs did not increase between 8-16 hours of BCG-CM treatment (Figure 9A). Moreover, we did the co-IP assay of ABCG2+ CSCs which showed increase in HMGB1 bound p53 by 3-fold (160 +/- 22 ng/ml versus 485 +/- 32 ng/ml; n= 4; p = 0.02) between 8 and 16 hours of BCG-CM treatment (data not shown). This data further confirms the release of fresh HMGB1/p53 complex by the BCG-CM treated ABCG2+ CSCs. Importantly, the culture supernatant containing this high level of HMGB1/p53 complex further induced bystander apoptosis to another untreated population of ABCG2+ CSCs (data not shown), suggesting amplification of the original alarm/death signal.

**Figure 9:**
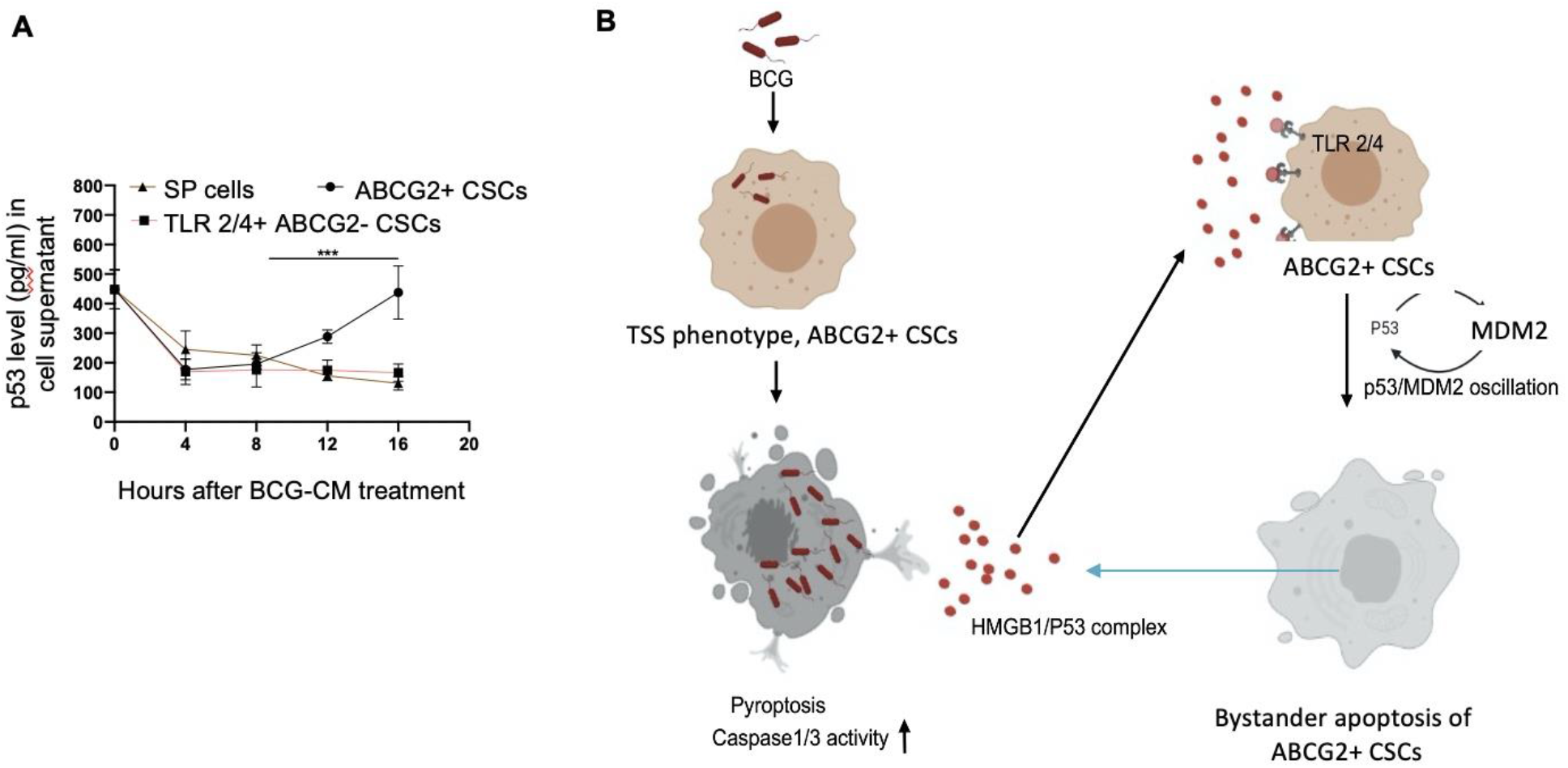
TSS phenotype can amplify the pathogen induced bystander apoptosis (PIBA). **A.** The p53 uptake assay in the culture supernatant was measured from 0-16 hours of BCG-CM treatment in the cells. The SCC-25 SP cells were obtained as described in figure 1. **B.** Potential mechanism of altruistic niche defense of CSCs against BCG infection. In the infected CSCs, as part of the ASC based niche defense mechanism [25], [40], [53], HMGB1 form a complex with cytoplasmic p53 to make an, “altruistic cell death signal”. The nature of the HMGB1 and p53 binding is not yet clear, but may form a stable complex as previously shown [30]. The TLR4, which is known to participate in receptor-mediated endocytosis, and possibly TLR2, internalizes the HMGB1/p53 complex leading to the induction of altruistic cell death signaling characterized by p53/MDM2 oscillation and activation of p53-induced pro-apoptotic genes. The ABCG2+ CSCs undergoing bystander apoptosis releases the HMGB1/p53 death complex, amplifying the altruistic death signal. This entire mechanism can be considered as pathogen induced bystander apoptosis (PIBA) and innate defense mechanism of stem cell niche. Although, in vitro, PIBA target only the TSS phenotype, in vivo, PI-BA may target the other cancer cell phenotype, as demonstrated by the marked anti-tumor activity of BCG-CM [Figure 3D].

## Discussion

Cancer stem cells (CSCs) promote invasion, metastasis, and drug resistance. CSCs may reside in the hypoxic niche, and reprogram to a highly aggressive phenotype; tumor stemness switch (TSS) phenotype. Therefore, targeting these CSCs in the hypoxic niche with the existing therapeutic strategies is clinically challenging. Here, we demonstrate that post hypoxia ABCG2+ CSCs of oral squamous cell carcinoma (OSCC) exhibit pyroptosis when infected with BCG. The CM of BCG-infected ABCG2+ CSCs is capable of inducing bystander apoptosis of non-infected CSCs by releasing a death signal, the HMGB1/p53 complex. This death signal then induces p53 mediated apoptosis of bystander cancer cells in a TLR 2 and 4 dependent manners. Thus, our work indicates that pathogen infected CSC of TSS phenotype releases death signal that eliminates the nearby CSCs. We suggest this form of pathogen-induced bystander apoptosis (PIBA) as a novel mechanism of TSS phenotype mediated “stem cell niche” defense mechanism that we recently described in virus infected MSCs [25].

PIBA is a part of the innate immune defense mechanism that protects host cells from invading pathogens [14]. Kelly DM *et al* reported *Mycobacteria* mediated PIBA in macrophages; where *Mtb* infected macrophages exert PIBA by direct cell to cell contact [15]. The HIV infected CD4+ T cells exert PIBA, which is mediated by the viral envelope protein [28]. We speculated that PIBA may be involved in stem cell niche defense mechanism, which we recently reported in virus-infected lung alveolar MSCs [25]. Thus, bystander apoptosis may represent a stem cell niche defense mechanism against pathogen invasion in the niche, whereby uninfected potential niche cells are eliminated to limit pathogen’s invasion. We speculated that like normal stem cell niche, CSCs may also exert a PIBA based niche defense mechanism, which may be exploited to eliminate CSCs in their niches.

BCG being an FDA-approved immunotherapy in invasive bladder cancer we wanted to explore a putative BCG-induced CSC niche defense mechanism. The *Mtb-m18b* strain that showed activation of stem cell niche defense by MSCs [25] served as positive control. Thus, the CSCs of several cell lines that we previously characterized for the TSS phenotype (SPm hox enriched in ABCG2+ CSCs) including oral cancer, breast cancer and lung cancer were infected with *Mtb-m18b* as well as BCG to evaluate PIBA. To evaluate if the pathogen can induce bystander apoptosis in CSCs without TSS phenotype, we used ABCG2-CSCs. We found that the pathogen replicated intracellular to ABCG2+ CSCs and the CM of these cells showed anti-bacterial activity as well as bystander cell death in freshly obtained ABCG2+ CSCs (Figure 1). Whereas, the pathogens did not infect and replicate intracellular to ABCG2-CSCs (Figure 2); CM of these infected cells did not induce bystander apoptosis of untreated corresponding cancer cell phenotype (Figure 1), suggesting that PIBA is limited to TSS phenotype only. Therefore, we used post-hypoxia/oxidative stress ABCG2+ CSCs of SCC-25 cell line to further investigate the molecular mechanism of PIBA. We found that the BCG-CM treated ABCG2+ CSCs was associated with significant up-regulation of caspase-1 and gasdermin D on day-12 (Figure 2). These results confirm that the pathogens selectively infect, replicate and then induce pyroptosis in the ABCG2+ CSCs versus ABCG2-CSCs. The conditioned media (CM) from the day-12 infected ABCG2+ CSCs activated a caspase1/3 mediated apoptosis in the freshly isolated ABCG2+ CSCs, confirming PIBA. Notably, the treatment of infected ABCG2+ CSCs with disulfiram (an inhibitor of pyroptosis) significantly reduced pyroptosis. Moreover, the CM-of disulfiram treated ABCG2+ CSCs failed to induce bystander apoptosis. These results suggested that during pathogen induced pyroptosis, some soluble factors were released, and that mediated bystander apoptosis. We identified the HMGB1/p53 complex as a soluble factor that mediated bystander apoptosis. Subsequent findings suggest that ABCG2+ CSCs under-going bystander apoptosis releases HMGB1/p53 complex into the culture supernatant, thus amplify the bystander apoptotic signal. Whereas, SP cells as well as ABCG2-CSCs undergoing bystander apoptosis failed to release HMGB1/p53 complex into the culture supernatant. This may explain the reason of low level of bystander apoptosis in this cell phenotype (Figure 1). We note that detailed molecular investigations are needed to confirm the mechanism of bystander apoptosis.

The HMGB1 has potent immuno-suppressive [29] and pro-survival properties as it activates nuclear factor kappa B (NF-kB) via TLR signalling pathways. However, HMGB1 may form complex with p53, and this complex can regulate apoptosis and autophagy in human colon cancer cell line HCT116 [26]. Whereas, HMGB1/p53 complex may mediate altruistic cell death in the human Embryonic Stem Cells (hESC) derived ASC phenotype characterized by the activation of p53/MDM2 oscillation [24]. We consider this altruistic cell death as an important component of the putative ASC based stem cell niche defense mechanism [24], [25]. In this context, our findings of BCG-CM mediated HMGB1/p53 release and the activation of p53/MDM2 oscillation may be viewed as a part of the ASC based stem cell niche defense mechanism being activated by TSS phenotype of ABCG2+ CSCs. Further investigation is required to find out whether the activation of ASC based stem cell niche defense mechanism could be exploited as a novel therapeutic strategy to target CSCs in their niches.

The TLR-2 and 4 may be involved in this putative TSS phenotype mediated niche defense mechanism as the inhibition of these two receptors significantly reduce bystander apoptosis of ABCG2+ CSCs. The TLRs are an integral part of innate immune defense mechanism. However, in cancer, TLRs act as a double edged sword in either favoring stemness [10], [31], [32] or apoptosis [33]. BCG activates TLR2/4 in foam macrophages and T cells to induce metabolic reprogramming [34], and T cell response [35]. In the bladder cancer cells, BCG was shown to induce apoptosis via TLR-7 [56]. However, in our study, we found the involvement of TLR2/4, but not TLR 7 in the PI-BA of ABCG2+ CSCs. TLR 2 and 4 are surface receptors, and their cellular signaling mechanisms are mediated primarily by MYD88 adaptor protein [31], [32]. TLR 4 also mediates endocytosis [48] and therefore, may be capable of internalizing the HMGB1/p53 complex. We propose that the HMGB1/p53 complex may internalize via TLR 4 and TLR 2 into the cytoplasm of CSCs, and then activate endogenous p53 for the induction of p53/MDM2 oscillation (Figure 9). Future studies are required to unravel the role of TLR2 and 4 in the internalization of HMGB1/p53 complex into the ABCG2+ CSCs and in the induction of p53/MDM2 oscillations.

During the last two decades, various strategies have been explored to activate p53 in tumor cells including small molecules that can disrupt MDM2, inhibit nuclear translocation, and or reverse mutant to wild type conformation. However, none of these mechanisms involve the induction of p53/MDM2 oscillation. An alternative mechanism could be the targeting molecular pathways that suppress p53 in CSCs. Using EU-MYC model of T-cell acute lymphoblastic leukaemia (T-ALL), we reported that SCA1+ CSCs use MYC-hypoxia Inducible Factor 2α (HIF-2α) stemness pathway to suppress p53 and decrease ROS production [1]. The MYC-HIF-2α stemness pathway is also active in the ABCG2+ CSCs of SCC-25 cell line [41]. In hESCs exhibiting altruistic behavior, the HIF-2α stemness pathway was activated and inhibition of the pathway led to the induction of p53/MDM2 oscillation [24]. Whether the BCG-CM mediated p53/MDM2 oscillation of ABCG2+ CSCs is the result of inhibiting the MYC-HIF-2α stemness pathway is now under active investigation.

It is critical to target the CSC population in a tumor. Recent developments in the field of CSC targeting therapeutics include several approaches such as targeting CSC surface markers [36], and targeting CSC signaling cascades like Notch, Hedgehog, Wnt, NF-κB [37] However, none of these strategies are showing encouraging results in clinical trials. Hence, the PIBA of CSCs may have a potential significance as a novel strategy to target CSCs.

In Summary, here we provide experimental evidences that the TSS phenotype of SCC-25 derived CSCs exhibit niche defense against BCG infection. We found that the CM of BCG-infected CSCs release HMGB1/p53 complex, which then induces p53/MDM2 oscillation in uninfected CSCs of TSS phenotype. This bystander apoptosis can be inhibited by neutralizing antibodies against HMGB1, TLR 2 and TLR 4, or small molecular inhibitor of p53. We speculate that the BCG-induced bystander apoptosis is a part of the recently identified ASC based stem cell niche defense mechanism against pathogen invasion. Understanding this TSS phenotype exhibiting CSC mediated niche defense may help to gain insight about tumor progression, as well as develop innovative strategies to target CSCs in their niches.

## Materials and methods

### Bacterial strains and Culture

All the necessary experimental procedures were undertaken inside BSC-class II facility in accordance with guidelines of “Institutional Bio-safety Committee” of KaviKrishna Laboratory. BCG strain (ATCC^®^ 35737™) was obtained from the American Type Culture Collection (ATCC) and grown in DifcoTM Middlebrook 7H9 broth (Becton Dickinson). The media contained 0.5% glycerol, 0.06% Tween-80, and 10% oleic acid albumin dextrose catalase (OADC), BD, USA). Streptomycin-auxotrophic mutant *Mtb strain18b* (gifted by Prof. Stewart T. Cole, Ecole Polytechinque Federale de Lausanne, Lausanne, Switzerland) was cultured in 7H9 medium (Difco, BD Biosciences, Franklin Lakes, NJ) supplemented with Middlebrook albumin-dextrose-catalase, 0.5% glycerol and 0.06% Tween 80 (Sigma, St. Louis, MO) until an OD of approximately 1 was obtained. We added 50μg/ml of streptomycin sulfate into the *Mtb-m18b* culture medium for bacterial growth [39]. GFP tagging procedure was performed as described earlier [39], [20]. The bacteria were prepared as single cell suspension in RPMI media as described [39] before being used to infect CSCs.

### Cancer Cell culture and sorting of CSCs with TSS phenotype

SKN-BE-2, HOS and RH4, H-146, LOVO, MCF-7, SCC-25 cell lines from American Type Culture Collection (ATCC CRL-1628) Manassas; VA were maintained as previously described [1], [19]. The SCC-25 cells were cultured in Dulbecco’s modified Eagle’s medium containing Ham’s F12 (DMEM F-12) in the ratio of 1:1. DMEM F-12 is enriched with 1.2g/L sodium bicarbonate, 2.5mM L-glutamine, 15mM HEPES and 0.5 mM sodium pyruvate (catalog no. 11330-057; GIBCO). The media was supplemented with 400ng/ml hydrocortisone (catalog no. H0888; SIGMA) and 10% fetal bovine serum (catalog no. 16000-044), used as Complete Isolation Media (CIM). The cells were maintained in a humified atmosphere of 5% CO_2_ at 37°C [19] and the other cell lines were also maintained as described previously [19]. Hypoxia/oxidative stress (<0.1% O_2_) was generated in a sealed container using BBL GasPak Plus anaerobic system enveloped with a palladium catalyst (Becton Dickinson, Cockeysville, MD) as described previously. To obtain TSS phenotype of CSCs, side population (SP) cells were flow cytometry sorted [19] and then exposed to 24 hours of hypoxia followed by 24 hours of re-oxygenation. Then, migratory SP (SPm) and non-migratory SP (SPn) cells were collected as previously described [19], [52]. The post hypoxia SPm cells or SPm (hox) cells exhibit TSS phenotype, and highly express ABCG2 [19], [52]. For the SCC-25 cell derived ABCG2+ and ABCG2-CSCs, the post hypoxia/oxidative stress treated cells were first subjected to immunomagnetic sorting for EpCAM+ cells by using EpCAM antibody (#ab 213500) conjugated with FITC by SiteClick antibody labeling kit. This EpCAM+ cells were then expanded for 7 days in spheroidal culture media (serum free culture containing 20ng/ml EGF and BFGF) as described previously [19]. The ABCG2+ CSCs were then immunomagnetically sorted by using the ABCG2 antibody (#ab 3380, Abcam), conjugated with PE by SiteClick antibody labeling kit as previously described [19]. For the immunomagnetic sorting, a PE sorting kit (#18554, Stem Cell Technologies, BC) was used. Noted that both ABCG2+ and ABCG2-CSCs of SCC-25 cell lines expressed CD44, LDH1 and CD133 equally [41].

### BCG and *Mtb-m18b* infection of ABCG2+ CSCs and collection of BCG-CM

The immunomagnetically sorted cells were cultured in vitro for 48 hours, and then treated with BCG or *Mtb*-m18b with MOI 5:1 as previously described including treatment with amikacin 200 µg/ml to kill extracellular bacteria [39]. The infected cancer cells were then washed twice with serum-free RPMI, and incubated in the appropriate cell culture media for the desired time at 37^°^C and 5% CO_2_. The CM was collected at desired time starting from day 10 by adding fresh serum free 1 ml DMEM per 1x10^5 cells for 48 hours, and filter sterilized with 0.2µm filter, concentrated with Centricon centrifugal filter units (EMD Millipore) to 10X to prepare 0.1 ml CM containing 100 ng/ml protein. The CM was then utilized to treat fresh ABCG2+ CSCs to evaluate bystander apoptosis. To collect rifampicin (RIF) treated CM, the day-9 infected cells were treated with 2µg/ml rifampicin for 3 days to kill intracellular bacteria as previously described [39].

### *In vivo* tumorigenicity assay

All the necessary experimental procedures were undertaken in accordance with approvals of Institutional Animal Ethics Committee of KaviKrishna Laboratory, and Gauhati University. To generate subcutaneous tumors, 1x10^5 ABCG2+ CSCs of SCC-25 cells were mixed with Matrigel 100 µl, and then injected subcutaneously to NOD/SCID mice following proper ethical permission as described. After 6 weeks, when the tumor reached 0.5 mm3 size, the animals were locally injected with concentrated CM/week (2 ml concentrated to 0.1 ml containing 0.5mg protein) into the tumor. The tumor size was measured with a caliper on a biweekly basis for 10 weeks and tumor volume was determined using the formula 0.5ab^2^, where b is the smaller of the two perpendicular diameters as described [42]. After 10 weeks, tumors were dissociated and single cell suspension was obtained to perform clonogenic assay and to evaluate the frequency of ABCG2+ CSCs.

### Clonogenic assay

The single cell suspension of dissociated tumors were freshly sorted via immunomagnetic sorting and 1x10^3^ EpCAM+/ABCG2+ or EpCAM+/ABCG2-CSCs were seeded in methylcellulose medium (Methocult M3134, Stem Cell Technologies) as described previously [1], [19]. The cells were seeded in 6 well plates, incubated at 37°C and 5% CO_2_. The colonies were counted after two weeks [1].

### Real-Time PCR (qPCR)

The real time PCR was performed as described previously using the TaqMan Gene expression assay [1]. The glyceraldehyde 3-phosphate dehydrogenase (GAPDH) was used as an endogenous control and RNA was quantified by the delta delta CT method using Q-Rex software version 1.1 (Rotor-Gene Q-Qiagen, New Delhi, India). The following TaqMan gene expression primers were used. Human: ABCG2 (Hs00184979_m1), TLR 2 (Hs02621280_s1), TLR 4 (Hs00152939_m1), TLR 7 (Hs01933259_s1), TLR 9 (Hs00370913_s1), p53 (Hs01034249_m1), p21 (Hs00355782_m1), PUMA (Hs00248075_m1), Bax (Hs00180269_m1), and GAPDH (Hs00266705_g1), MDM2 (Hs01066930_m1), HMGB1 (Hs01923466_g1).

### Pyroptosis assay

The pyroptosis of BCG infected ABCG2+ CSCs was evaluated by measuring cleaved caspase 1 level by ELISA, Caspase 1 activity and lactate dehydrogenase (LDH) release assay. The Caspase 1 activity assay was performed by using the Caspase 1 substrate Ac-YVAD-AFC (Cayman Chemical, An Arbor, Michigan, USA) as previously described [44], [45]. Briefly, cell lysate prepared for the Caspase-3/7 assay was mixed with Ac-YVAD-AFC, and after an hour the fluorescence signal of cleaved AFC was detected at 400 nm excitation and 505 nm emission using fluorescence spectrofluorometer (Agilent Varian Cary Eclipse, Hyderabad, India). For the LDH release assay, the BCG-CM was subjected to LDH measurement by the LDH-cytotoxicity assay kit (#ab65393, Abcam) as per manufacturer instruction with slight modifications. Briefly, 25 µl of BCG-CM was mixed with 25 µl of LDH assay reagent, and the assay reaction was stopped after 30 minutes by adding 25 µl of stop solution. The OD value at 450 nm was taken using iMark Microplate Absorbance Reader (Biorad, Gurgaon, India). Some of the assay results were confirmed by Decker method [57]. To further confirm pyroptosis, LDH was also measured after treating the cells with disulfiram 50 nM/twice daily or Caspase 1 inhibitor z-YVAD-fmk.

### Cellular apoptosis or caspase-3/7 activity assay

The assay was performed as described previously by using the caspase-3/7 substrate Ac-DEVD-AMC [51], [45], [19], [1]. Briefly, 100 µg/ml of cell lysate was prepared by lysing 5x10^3 -1x10^5 cells using a modified 1X RIPA Buffer: Tris-HCl (20mM; pH 7.5), NaCl (155 mM), 1 mM Na_2_ EDTA (1 mM EDTA from 100 mM stock solution in H2O, pH 7.4), EGTA (1.5 mM EGTA), Triton (1.2%), sodium Pyrophosphate (25 mM), Sodium Fluoride (25 mM), β-glycerophosphate (1 mM), activated sodium orthovanadate (Na_3_VO_4_) 1 mM (from 200 mM stock solution), 1 µg/ml leupeptin, 1µg/ml aprotinin, and 1 µg/ml pepstatin. 1 mM Phenylmethylsulfonyl fluoride (PMSF) (200 mM stock solution prepared in isopropanol and stored at RT) and 5mM dithiothreitol (DTT) was added immediately before use. Cells in a 1.5 ml microcentrifuge tube were centrifuged in ice-cold phosphate buffered saline (PBS), and the pellet was treated with 50 µl of RIPA buffer, kept on ice for 10 minutes, and stored at -80°C. After a few days, lysate was thawed on ice, equal volume of freshly prepared RIPA buffer was added, vortexed for 1 minute, and kept on ice for 5 minutes, then centrifuged at 4C/5000 RPM, and the supernatant was transferred to a fresh tube as 25 µl aliquot and stored at -80°C for future use. To perform the assay, a 25 µl aliquot was thawed on ice, and 200 µl of Ac-DEVD-AMC (Cayman Chemical, An Arbor, Michigan, USA) substrate (prepared by adding 0.1 ml of 1mg Ac-DEVD-AMC in DMSO to 4 ml of the lysis buffer containing freshly added DTT and PMSF) was added on to it. The mixture was vortexed, and the enzymatic activity was measured by detecting cleaved substrate linked to fluoropore using a fluorescence spectrofluorometer (Agilent Varian Cary Eclipse, Hyderabad, India) as described previously [45].

### Caspase inhibition

The experiment was performed as previously described [1], [19], [45]. Anti-Caspase 8 is the Z-IETD-FMK (Fluoromethyl ketone); R&D Systems #FMK007 and Anti-Caspase 9 is Z-LEHD-FMK (#FMK 008). For both caspase inhibition experiment, 100 uM (dissolved in DMSO) of each was added in the cell culture for 4 days. The medium was changed every second day.

### Inhibition of ferroptosis, necroptosis and autophagy

Ferrostatin-1 (20 uM), necrostatin or RIP-I kinase (20 uM), and 3-Methyladenine (3-MA) of 5 mM was prepared in DMSO (from 100 mM stock). These reagent mixtures were used to inhibit ferroptosis, necroptosis and autophagy respectively. Reagents were obtained from Sigma-Aldrich; 2 μM (in vitro) and neutralizing HMGB1 of 10µg/ml were obtained from Biolegend (isotype control mouse IgG2a kappa, #16-4724, Biolegend.

### Enzyme Linked Immunosorbent Assay (ELISA)

The cell lysates were prepared by RIPA buffer and subjected to ELISA assay as described [45], [25]. The information about various ELISA kit and antibody details is given in supplementary method, Table 1. The absorbance was measured at 450 nm using iMark Microplate Absorbance Reader (Biorad, Gurgaon, India).

**Table 1:** The information about various ELISA kit and antibody details

| Protein name | Antibody | ELISA kit |
| --- | --- | --- |
| p53 | 2527 (Cell Signaling Technology) | DYC1043-2 R&D |
| MDM2 |  | DYC1244-2, R&D |
| Beta-Actin | #3700 (Cell Signaling Technology) |  |
| Vinculin | #4650 (Cell Signaling Technology) |  |
| TLR 4 | NB100-56566 (Novus Biologicals) for ELISA/WB.<br>Mabg-htlr4 (InvivoGen, San Diego, CA) for neutralizing. |  |
| Cleaved Gasdermin D (GSDMD) | #36425 rabbit polyclonal (Cell Signaling Technology) and<br>#H00079792-M01 (Abnova). |  |
| TLR2 | NB100-56722 (Novus Biologicals) for ELISA/WB.<br>Maba2-htlr2 (InvivoGen, San Diego, CA) for neutralizing. |  |
| HMGB1 | H00003146-M08 (Novus Biologicals, Littleton, CO) Western blot, and<br>neutralization.<br>#ab228624 A (Abcam) for IP. | #NBP2-62766, Novus Biologicals |
| Cleaved caspase 1 | #A1004 rabbit polyclonal (Biovision, Milpitas, CA).<br>#sc-56036 mouse monoclonal (Santacruz) |  |
| ABCG2 | AB3380 (Abcam) | MBS703358<br>Mybiosource, CA |
| Cleaved caspase 3 | PAS-114687 rabbit polyclonal (Thermofisher scientific). | DYC835-2 |

### Western blot and co-immunoprecipitation

Western blot analysis was done on a 4-12% sodium dodecyl sulfate–polyacrylamide gel electrophoresis (SDS-PAGE) gel and transferred to polyvinylidene difluoride (PVDF) membranes (Millipore-Sigma, Immobiolon-P, Cat # IPVH20200) as previously described [1], [19]. Co-immunoprecipitation of the HMGB1-p53 complex in the BCG-CM was performed following lab’s standard IP protocol [45], [1] using the protein A sepharose beads (GE Cat # 17-0780-01; Millipore-Sigma). The concentrated BCG-CM (containing 1.0 mg protein in the lysis buffer) was subjected to cross-linking by DTSSP (Thermofisher Scientific #21578) as per manufacturer instructions before performing IP. Prior to IP, samples were pre-cleared with Protein A Sepharose at 4^0^C for 3 hours with gentle shaking. A rabbit polyclonal antibody (10 ug of #ab 228624, Abcam) was used to allow HMGB1 immune complex to form (in 500 ul solution containing 1.0 mg protein), which was captured with 50% slurry of BSA blocked Protein A Sepharose beads. The immune complex was then eluted by boiling the beads in 2X SDS sample buffer, and the elutes were washed 4 times with 1X PBS with 0.2% Tween 20. The elutes were subjected to WB probing with a mouse monoclonal antibody (#H00003146-M08; Novus Biologicals) against human HMGB1. To confirm the p53 IP, blots were stripped (Thermo Fisher Restore stripping buffer, Cat # 21059) and re-probed using p53 antibody (#2527, Cell Signaling Technology). Inputs representing 2.5% of the lysate subjected to immunoprecipitation was further subjected to WB for HMGB1 using the mouse monoclonal antibody. The Co-IP elutes (eluted with elution buffer containing glycine and Tris-HCL and 500 mM NACL) were also subjected to ELISA to quantify p53 and HMGB1 proteins after the elutes were neutralized by 10X PBS. Isotype control with a rabbit polyclonal IgG (ab #37415; Abcam) was run in parallel to the sample in each IP procedure.

### Silencing of p53

The inhibition of p53 was achieved by Accell siRNA (Thermoscientific Dharmacon, Lafayette, CO, USA,) and by pifithrin-α, an inhibitor of p53 as described previously [19]. The Accell siRNA used for p53 was A-003329-22-0005. Briefly, the ABCG2+ CSCs (10^4^ cells/well in 96-well plate) before treating with BCG-CM were treated with 1 μM Accell siRNA as per manufacturer instructions. After 72 hours of incubation at 37°C, gene silencing was confirmed using real time qPCR.

### Antibody blocking experiments

The experiment was performed as previously described [1], [19], [45]. Anti-Fas monoclonal antibody (human, neutralizing) clone ZB4 (Sigma-Aldrich); Anti-human TNF-monoclonal antibody (MAB 210;R&D Systems, Minneapolis, MN); Anti-human TRAIL (clone RIK-2, Thermofisher Scientific, Waltham, MA);Anti-hLAP or TGF beta 1 (MAB 246; R&D Systems, Minneapolis, MN);Anti-TLR2 (cloneTL2.1; BioLegend, San Diego, CA), and Anti-TLR4 (HTA125; BioLegend, San Diego, CA); Immunoglobulin G1 (IgG1) isotype control antibodies were used at corresponding concentrations. Antibodies were added to BCG-CM, mixed well before adding to ABCG2+ CSC grown in 6-well culture plates. The abilities of the blocking antibodies to neutralize their ligands were determined by challenging Jurkat or undifferentiated THP-1 cells with the appropriate ligand in the presence of the antibody at the concentrations indicated above. Fas ligand (MAB050), TRAIL (375-TL-010), TGF-beta 1 (7754-BH-005) and TNF-alpha (210-TA) were obtained from R&D Systems, and the synthetic bacterial lipopeptide Pam3CysSerLys4 was obtained from Calbiochem. The cells were incubated with cycloheximide for 15 min before addition of the apoptotic stimulus.

### Statistical Analysis

The statistical calculations were performed using either Student’s t test or One-Way ANOVA with Dunnet *post-hoc* test by GraphPad Prism version 8.4.2). Data are expressed as means ± SEM; *\*p<0.05, **p<0.01,*** p<0.001, ****p<0.0001*.

## Acknowledgement

We thank Dr. Antonio Campos Neto and Dr. Philip Stashenko, Forsyth Institute, Cambridge for their valuable suggestions in this research work. We also thank Dr. Jyotirmoi Phukan, Gauhati Medical College and hospital and Dr. Anupam Sarma, B. Borooah Cancer Institute for their valuable suggestions. We thank Mr. Biswajit Das and Mallika Maral for taking care of the animal facility. We thank the members of KaviKrishna Laboratory, Indian Institute of Technology Guwahati Research Park, Guwahati, Assam, India, Thoreau Laboratory for Global Health, M2D2, University of Massachusetts, Lowell, Massachusetts, and Department of Bioengineering and Technology, Gauhati University, Guwahati, Assam, India.

## Author contributions

B.D. initiated and designed the study. B.D., S.B., B.P., L.P., P. J. S., S. M., S.G., R. B. M., and H.L performed the in vitro and in vivo experiments. B.D., S.B., B.P., and L.P. analyzed data. B.D., S.B., L.P., and S.G. wrote the article. B.D., L.P., P.J.S., S.M., C.V.R., and D.B. edited the article.

